# A network theory of the origin of life

**DOI:** 10.1101/096701

**Authors:** Geoffrey W. Hoffmann

## Abstract

The origin of life may have been a low probability event that involved both RNA and polypeptides. While such an event would occur with low probability, the probability is not too low in the context of the number of nucleation opportunities on the primitive Earth. The probability for the nucleation event is too small if needed components occur as frequently as disruptive components do, without specific interactions between the two classes. However, if many of the needed components contribute by inhibiting otherwise disruptive components, a high probability for a complex nucleation as the beginning of life event emerges. RNA interference and long non-coding RNAs are ascribed roles in inhibiting components that would otherwise be disruptive in the context of a nucleation event. The theory likewise provides an inhibitory role for the large fraction of proteins (35% to 40%) that have no known function in both prokaryotes and eukaryotes. It is suggested that volcanic ash could provide the large number of random shapes needed as the basis for the nucleation event. Experiments based on that hypothesis are proposed.

---

The problem of finding an explanation for the transition from inanimate matter to life is a significant challenge for science. The simplest living matter we know is already complex. For example, the bacterium E. coli is one of the simplest autonomous living organisms, and it has 4,288 genes. These genes are transcribed to produce essential ribosomal RNA, transfer RNA and thousands of messenger RNA molecules that are translated to produce thousands of proteins. In light of the complexity of life as we know it, the idea that it could have started as an improbable nucleation event involving both replication and translation of RNA, and going suddenly from non-living to living matter, is not usually considered. We here present a model that supports such a complex nucleation event as the origin of life. The envisaged nucleation event involves both “plan” molecules in the form of nucleic acid, and polypeptides that are “tools” encoded in the nucleic acid.

We present a calculation of how complex a nucleation event could be in the context of the time and space available on the primitive Earth. We begin by estimating the number of opportunities there were for the nucleation event. This is the product of (a) the number of time elements between the formation of the earth and the time of the earliest fossils, and (b) the number of volume elements, for example in the oceans. It is the logarithm of this product that is important, and the level of complexity of the nucleation event is surprisingly insensitive to the exact assumptions regarding the sizes of the time and volume elements. We envisage life starting at one place and one time, in a volume element that contained a number *α* of needed components, in the absence of a number *β* of components that would be disruptive. We calculate this joint probability for a volume element of optimal size. In our model the joint probability is a function of only *a* and *β*.

We consider for example a volume element ∆V the size of a eukaryotic cell of size 10 × 10 × 10 cubic microns, that is, 10^−15^m^3^. This is for illustration only; this may not be the optimal volume element size. We further consider the case that the nucleation event occurred in the primitive oceans, where there may have been vast amounts of volcanic ash, which would provide random shapes, some of which would have been useful catalytic activities, some of which would have been disruptive catalytic activities (in the context of a given set of needed activities), and the majority of which would have had no relevant catalytic activity at all. The total volume V of the oceans is approximately 10^17.8^m^3^, so the number of volume elements that are available, namely V/∆V, could be up to of the order of 10^33^.

Let ∆T be a time element sufficient for the production of a rudimentary set of needed components within the volume element that is of optimal size for containing a set of needed components and no disruptive components. Then the number of time elements available is T/∆T, where T is the time from the formation of the Earth (4.5 billion years ago) to the time of appearance of the oldest known fossils (3.5 billion years ago), that is, one billion years or 3 × 10^16^ seconds. For illustration I consider the case that the size of a time element ∆T is one day, or 86,400 seconds. Then TMT is approximately 2 × 10^10^.

An upper limit for the number of opportunities Ω for the origin of life within the space and time available could then be a very large number, namely

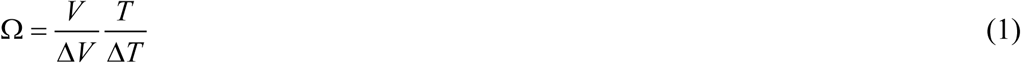

and for the example given, Ω is approximately 2 × 10^43^.

If *p* is the probability that any one of *N* random shapes present in a volume element has a catalytic activity, the probability *P* that the volume element contains one or more of each of *α* needed components and none of any of *β* disruptive components is given by

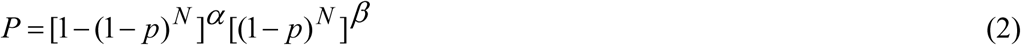

As *N* increases, *P* passes through a sharp maximum, denoted *P*_max_, located at 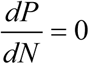, the magnitude of which is independent of *p*, and is a function of only *α* and *β*. For a volume element that is not too big and not too small, but just right for given values of *α* and *β*, we obtain^1^

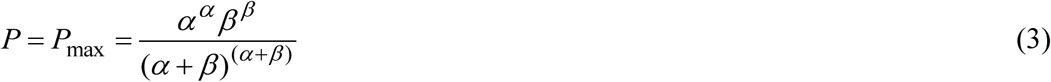

For the origin of life to be a probable event in the context of this model we need the product of *P*_max_ (a very small number) and Ω (a very large number) to be greater than 1.

For this model to be applicable we need volume elements of the optimal size. This is most readily the case if the nucleation event does not involve boundaries that would be an additional constraint on the size of the volume element. We consider first the simple case *α* = *β*, meaning needed activities occur as frequently as disruptive activities, without any interactions between the two classes. Then

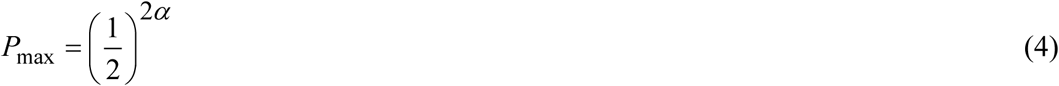

and for *P*_max_ Ω > 1 with Ω = 2 × 10^43^ we obtain *α* < 72. Most biochemists would however agree that 72 catalytic activities are too few for a functioning living entity.

We obtain much larger values of *P*_max_ if we break the symmetry between *α* and *β*. This leads to the possibility of a much more complex nucleation event. Specifically, we consider the case that a *α ≫ β*, and rationalize this version of the model by including among the *α* needed components activities that inhibit disruptive components. The inhibition can be at the level of nucleic acid-nucleic acid interactions, polypeptide-polypeptide interactions and polypeptide-nucleic acid interactions. Most of the disruptive activities present can then be inhibited, and it turns out that the number of needed components consistent with the time and space available can be surprisingly large.

Nucleic acid-nucleic acid inhibitory interactions exist in present day organisms as RNA interference^2^, including in the forms of microRNA^3^ and small interfering RNA^4^. RNA-RNA inhibitory activity may also include specific ribonuclease activities exerted by ribozymes. Most significantly, this model provides a role for long non-coding RNAs (lncRNAs), the function of which has been a mystery^5^ These RNAs are a highly conserved through evolution. One lncRNA has a structure that looks like 16S RNA, a highly conserved molecular machine.

Inhibitory polypeptide-polypeptide interactions could include blocking of catalytic sites and specific proteolysis of undesired polypeptides. Inhibitory polypeptide-nucleic acid interactions could include RNA sequence-specific ribonuclease activity and polypeptide-specific peptidase activity exerted by ribozymes.

Then for *α ≫ β*

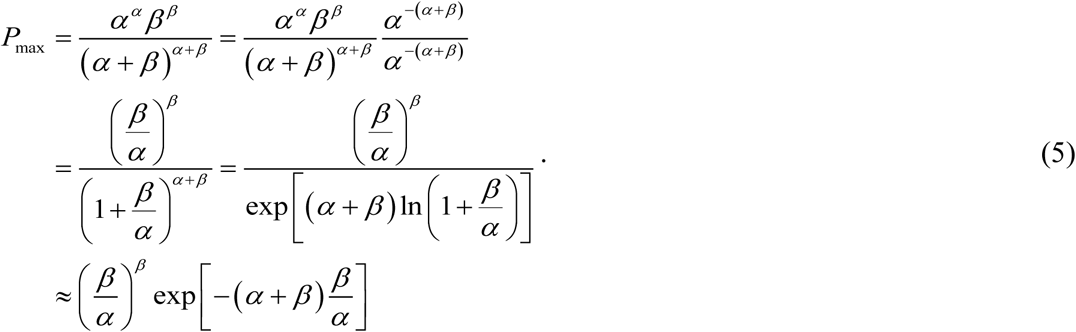

The final approximation holds because for *α ≫ β*, we can drop all but the first term in the Maclaurin expansion of ln (1 + *x*) = *x* − *x*^2^ / 2 + *x*^3^ / 3 −… to get

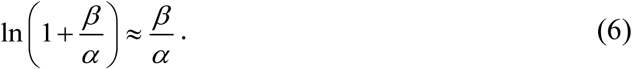

Continuing to approximate the final factor, we have

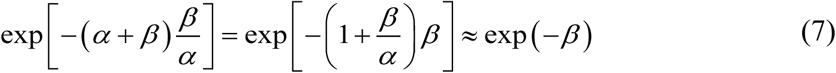

because *α ≫ β* permits us to drop the second term within the exponential. From the last line in equation (5), we have

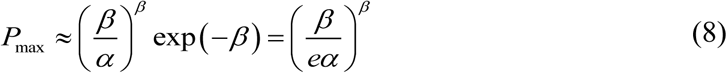

Hence we see that while *P*_max_ decreases exponentially as a function of *α* and *β* when *a = β* (equation 4), it decreases much more slowly in (8), namely as a polynomial function of *α*, when *α ≫ β*. For the latter case, examples of the values of *P*_max_ are as follows:

| $\alpha$ | $\beta$ | $\log_{10} P_{\max}$ |
| --- | --- | --- |
| 100 | 10 | -14.6 |
| 1000 | 10 | -24.4 |
| 10,000 | 10 | -34.3 |
| 50,000 | 10 | -41.3 |

Hence in this model, for *α* = 10,000 and *β* = 10, for *P*_max_ Ω > 1, log10 Ω can be as little as 34.3. The 10,000 needed activities could for example include about 5,000 activities that inhibit disruptive activities and 5,000 that are actually used in a positive way. It follows that in this model the nucleation of life would have an even chance of occurring in a total volume V that is much smaller than was assumed in our calculation of *P*_max_, namely as little as one billionth of the volume of the oceans.

The ribosome has a central role in translation. In prokaryotes it consists of three rRNA molecules and fifty-two proteins. While most disruptive activities can be inhibited by a complementary sequence (in the case of disruptive RNAs) or a complementary shape (in the case of disruptive proteins), this may not be the case for RNA that is complementary to rRNA, which we could call “anti-rRNA”. The anti-rRNA would be disruptive by inhibiting rRNA, and could in principle be inhibited by RNA that is complementary to anti-rRNA (“anti-anti-rRNA”), but this anti-anti-rRNA can itself be expected to be disruptive, because while similar but not identical to the rRNA, it could be expected to contribute to the formation of pseudo-ribosomes that are not functional. This would come with a high metabolic cost. Hence rRNA is a special class of RNA, for which inhibition of incorrect sequences is not readily achieved. It is partly on this basis that we have chosen *β* = 10 in the above example. The nucleation event may have involved more than three primordial rRNA molecules, with the joining of the separate rRNA molecules to yield just three as a subsequent step. If we choose *β* = 3, the probabilities obtained are even larger than for *β* = 10.

A plausible sequence of events then involves two steps. For example, a volume element ∆V contains 2,500 catalytic activities in the form of volcanic ash that suffice to get the system started, including the production of nucleotides, amino acids, amino acyl synthetases, ribosomal RNA and proteins, an RNA polymerase and a set of tRNAs. Then mRNAs are produced that encode a further 2,500 components, with a similar or equivalent set of functionalities. Thus the entire system nucleates with an RNA genome and a high probability in the context of the time and space available. A model calculation shows stable translation can involve an initial set of codon-amino acid adaptors with low fidelity^6^, which adds plausibility to this scenario.

The idea that life began with a system involving both polypeptides and nucleic acids was proposed in 1971 by Manfred Eigen in his hypercycle model^7^. He pointed out that the origin of life did not need to be as complex as is admissible in the context of the above calculation. For example, the nucleation event may have involved only the codons A and U, which allows for the involvement of diverse amino acids, including positively charged, negatively charged, hydrophobic and hydrophilic^7^, without the full complexity of the current genetic code.

This is a “frozen accident” theory for the origin of the genetic code. Both components that are needed in a positive sense and components that are needed in a negative (inhibitory) role would be part of the frozen accident. The need for both classes would then persist, and hence the theory provides roles for the many proteins that have unknown functions. More than 35% of proteins in prokaryotes^8^ and 40% of proteins in eukaryotes^9^ have unknown functions. The functional characterization of these proteins has been called “one of the main challenges of modern biology” ^9^ For a particular set of needed components we need a set of inhibitory proteins that complements the needed set. Many of the proteins of unknown function are associated with ribosome assembly, photosynthesis and cell wall pathways^9^. Hence in addition to finding a plausible model for the origin of life, we appear to have found a solution to the problem of proteins of unknown function.

Most significantly, this theory suggests new experiments that would enable it to be either disproven or developed in greater quantitative detail. For example, a racemic mixture of prebiotic molecules in a suitable abiotic catalytically noisy environment such as an aqueous suspension of ash is expected to be able to produce local asymmetry that can be detected optically. In small experiments a detector that scans a sample for the presence of such local asymmetry could be constructed. A suitable volume could be scanned for localized production of various metabolites, for example the production of particular amino acids or nucleotides. The production of more complex metabolites from less complex metabolites may be detected. This could include for example scanning the sample for the production of ATP from suitable precursors. It may be possible to detect and quantitate the production of polynucleotides from oligonucleotides and the replication of polynucleotides. The detection of functioning ribosome-like particles starting from an incomplete set of ribosome components may be possible. Eigen called this idea a “minus one” experiment^7^.

Networks are a recurring theme in biology. In addition to gene and protein networks this includes the immune system network and neural networks. Counterintuitively, life may have begun as a complex network.

## References

1. Hoffmann, G. W. The stochastic theory of the origin of the genetic code. Ann. Rev. Phys. Chem. 26, 123–144 (1975).

2. Fire, A. et al. Potent and specific genetic interference by double-stranded RNA in Caenorhabditis elegans. Nature 391, 806–811 (1998).

3. Chen, K. & Rajewsky N. The evolution of gene regulation by transcription factors and microRNAs. Nature Rev. Genetics 8, 93–103 (2007).

4. Hamilton, A. J. & Baulcombe D. C. A species of small antisense RNA in posttranscriptional gene silencing in plants. Science 286, 950–952 (1999).

5. Chi, K. R. Finding function in mystery transcripts. Nature 529, 423–425 (2016).

6. Hoffmann, G. W. On the origin of the genetic code and the stability of the translation apparatus. J. Mol. Biol. 86, 349–362 (1974).

7. Eigen, M. Selforganization of matter and evolution of biological macromolecules. Naturwissenschaften 58, 465–523 (1971).

8. Doerks, T., von Mering, C. & Bork, P. Functional clues for hypothetical proteins based on genomic context analysis in prokaryotes. Nucl. Acids Res. 32, 6321–6326 (2004).

9. Horan, K. et al. Annotating genes of known and unknown function by large-scale coexpression analysis Plant Physiol. 147, 41–57 (2008).

